# Diversity and prevalence of sulfate- and sulfite-reducing microorganisms in the human gut microbiome

**DOI:** 10.1101/2025.10.20.683523

**Authors:** Rebecca A. Christensen, Albert L. Müller, Alfred M. Spormann, Jonas B. Cremer, Jessica A. Grembi

## Abstract

Microorganisms capable of producing hydrogen sulfide have been associated with inflammatory diseases within the human gastrointestinal tract. Using a weighted protein homology search followed by phylogenetic fragment insertion, we sought to characterize the diversity and prevalence of microbial dissimilatory sulfite reduction genes (*dsrAB*) from over 70,000 human gut samples from publicly available metagenomic sequencing data. We found over 1,700 unique *dsrAB* sequences that were predominantly within two clades of sulfate- and sulfite-reducing microorganisms (SSRM): the well-studied Desulfovibrionaceae family and the Firmicutes group *sensu lato* (s.l.). We detected SSRM in over a third of adults, but very few children. *Bilophila wadsworthia* was the most prevalent SSRM, and along with other members of Desulfovibrionaceae, constituted the majority of SSRM found. About one fifth of people with SSRM contained members of the Firmicutes group s.l. Our study characterizes the overall diversity of SSRM in the human gut from different global populations while highlighting the potential importance of SSRM from the Firmicutes group, which have commonly been overlooked as contributors to microbial hydrogen sulfide production in the human gut.

## Introduction

Microbes in the human intestine play crucial roles in host health with both beneficial and detrimental effects (Clemente et al., 2012). The complex relationship between the host and the microbiota is mediated in part by the different metabolites which microbes produce. One such microbial metabolite is hydrogen sulfide (H_2_S) which has been suggested to severely affect gastrointestinal physiology and health through multiple mechanisms. H_2_S can compromise the integrity of the protective gut mucosa by breaking down the mucus layer and exposing the gut epithelium to harmful toxins and pathogens present in the gut lumen (Ijssennagger et al., 2016); disrupting colonic epithelial cell function by inhibiting the utilization of butyrate, a major microbial fermentation product that is an energy source for epithelial cells (Beaumont et al., 2016); and possessing tumorigenic and genotoxic effects even at moderate physiological concentrations (Attene-Ramos et al., 2006, 2010; Untereiner et al., 2017)). In line with these findings, H_2_S production has been associated with diseases characterized by gut inflammation, such as irritable bowel syndrome, ulcerative colitis, Crohn’s disease, and colorectal cancer (Dordević et al., 2020; Guo et al., 2016; Kushkevych et al., 2019; Lin et al., 2023; Singh & Lin, 2015; Wang et al., 2021).

Microbes in the gut can produce H_2_S when metabolizing a variety of sulfur-containing substrates, including sulfate, sulfite, thiosulfate, taurine, cysteine, and sulfoquinovose (Hanson et al., 2021; Laue et al., 1997; Müller et al., 2015). Sulfates and sulfites are of particular note as they can be used as electron acceptors by sulfate- and sulfite-reducing microbes (SSRM), allowing for anaerobic respiration where H_2_S is released in the final reaction step (Carbonero et al., 2012). While SSRM are a phylogenetically diverse group of prokaryotes, previous research on SSRM in the human gut has focused on the prevalence of a handful of well known species, namely *Bilophila wadsworthia* and *Desulfovibrio piger* (Fite et al., 2004; Gibson, Cummings, et al., 1988; Gibson, Macfarlane, et al., 1988; Zinkevich & Beech, 2000). Studies have further revealed a correlation between presence of these species and gut inflammation, suggesting that H_2_S production via sulfate/sulfite reduction can have detrimental health effects (Coutinho et al., 2017; Z. Feng et al., 2017; Figliuolo et al., 2017, 2018; Jia et al., 2012; Kushkevych et al., 2019; Loubinoux et al., 2002; Pitcher, 2000; Rey et al., 2013). However, these analyses mainly employed culturing approaches or PCR targeting of known gut-SSRM that might overlook many species of SSRM from other genera or even phyla. Studies in a variety of other environments have indeed shown that SSRM are found among many phyla and genera (Anantharaman et al., 2018; Diao et al., 2023; Müller et al., 2015; Pelikan et al., 2016; Wolf et al., 2022), which raises the question about the true diversity of SSRM in the human gut.

In this study, we aimed to characterize the prevalence and diversity of SSRM in the human gut by identifying SSRM from publicly available metagenomic sequencing data for 72,501 human gut samples. Previous research has phylogenetically characterized SSRM in the human gut by identifying sulfate reduction genes from metagenome assembled genomes (MAGs) of the human gut, highlighting the prevalence of relatively higher-abundance SSRM (Wolf et al., 2022). Here, we leveraged a computational approach to search partially assembled human gut metagenomes for the SSRM-specific dissimilatory sulfite reductase gene *dsrAB*, essential for catalyzing the final H_2_S-producing step of sulfite and sulfate reduction. Our approach 1) uses a highly curated database of all known *dsrAB* sequences; 2) allows for the inclusion of organisms difficult to assemble for various reasons (e.g. genomic repeats, low abundance, etc.) (Ghurye et al., 2016); an 3) includes samples across many geographic populations which are primarily not case-control studies of individuals with known gut inflammation. Using this approach, we identified that species of SSRM from the human gut fall primarily within two distinct clades of the *dsrAB* phylogenetic tree: the Desulfovibrionaceae family and the Firmicutes group *sensu lato* (herein referred to as the “Firmicutes group”). Furthermore, we found that nearly 40% of adults have gut microbes capable of microbial dissimilatory sulfite reduction.

## Results

### Construction and optimization of dsrAB profile HMMs

SSRM phylogeny and taxonomy can be derived from *dsrAB* sequence diversity as has been shown previously (Müller et al., 2015). To identify *dsrAB* sequences in the metagenomics datasets, we used HMMER, a position-dependent algorithm that weights alignment scores of target sequences based on their homology to the most highly conserved regions of a set of known sequences (see Methods). We used a highly curated set of 1,292 *dsrAB* reference sequences from a broad range of microbes (Müller et al., 2015) to construct a bacterial-type and an archaeal-type *dsrAB* profile Hidden Markov Model (pHMM) (see Methods). To validate the efficacy and sensitivity of these pHMMs in recovering *dsrAB* sequences, we further performed a HMMER search against the UniProt protein domain family containing sulfite reductases and the closely related nitrite reductase 4Fe-4S domain family (PF01077). PF01077 contains 34,408 protein sequences, including 541 *dsrAB* sequences and 33,876 non-*dsrAB* sequences. To reduce false positive rate below 0.25%, we chose a conservative bit score cutoff of 100 which resulted in the identification of >77 % of all annotated *dsrAB* sequences from PF01007.

### Diversity of SSRM inhabiting the gut

In order to identify SSRM species from the human gut, we mined publicly-available metagenomic sequencing data from 72,501 biosamples from the human gut available from the NCBI sequencing read archive (**Supplementary Table S1** includes the list of biosample IDs). A HMMER search using the bacterial-type *dsrAB* pHMM found 6,289 putative *dsrAB* sequences constituting 1,769 unique *dsrAB* sequences (herein referred to as “hits”). No hits were returned using the archaeal-type *dsrAB* profile. In order to determine the phylogeny of the SSRM, we placed the hit sequences into a *dsrAB* reference tree (Müller et al., 2015) via phylogenetic fragment insertion (**Supplementary Fig. 1**, see Methods). Hit sequences were assigned to the closest reference leaf. These putative *dsrAB* hits were most closely related to 39 reference leaves with 94% of hit sequences belonging to either the Desulfovibrionaceae family (71.9%) or the Firmicutes group (22.1%), see **Fig. 1A**. In order to evaluate SSRM diversity at approximate species level, we clustered these 39 reference sequences into 27 SSRM species-level clusters with >90% *dsrAB* nucleotide sequence identity (**Supplementary Fig. 2, Supplementary Table S2**), which corresponds to 99% 16S rRNA gene sequence identity (Müller et al., 2015).

**Figure 1.**
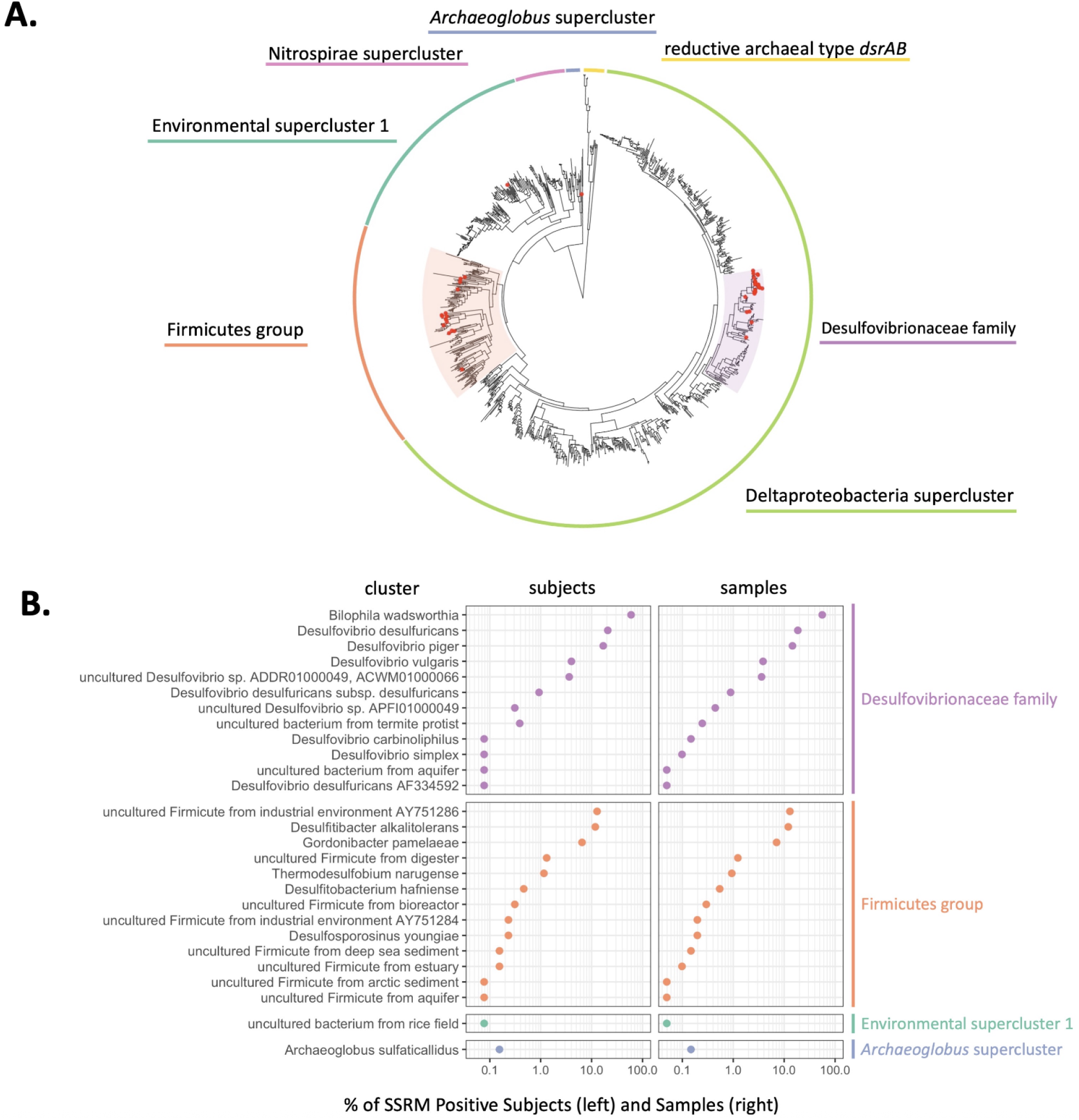
Diversity of SSRM in the human gut. A) Phylogenetic tree of reference *dsrAB* sequences. Arc color indicates major superclusters defined in Müller et al (2015). Reference leaves that were closest by branch distance to unique hit sequences found in this study from the human gut are indicated by red dots. B) Prevalence of each SSRM cluster in SSRM-positive biosamples and subjects. Clusters correspond to >90% *dsrAB* nucleotide sequence identity.

**Figure 2.**
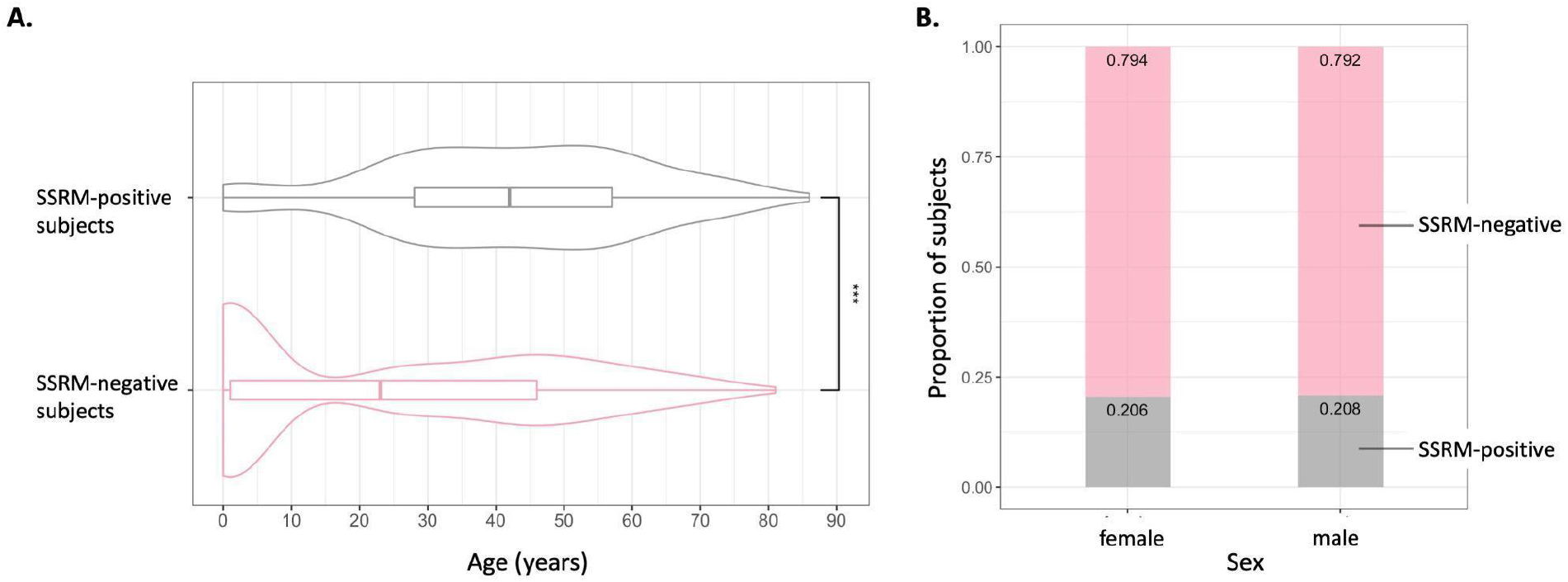
SSRM presence by age and sex. A) Subject age distribution by presence of SSRM. Significance of difference (***) confirmed by one-sided Wilcoxon rank sum test, p-value = 1. 63 * 10^−21^. B) SSRM presence by sex. No significant difference is detected (Fisher’s exact test, p-value = 1). Analyses based on 907 subjects with data on age and 179 subjects with data on sex (107 females, 72 males).

To determine SSRM cluster prevalence on the host level, we used available subject identifiers from the biosample metadata. Of the 72,501 biosamples, there were unique identifiers for 4,228 subjects, some of whom provided multiple samples (**Supplementary Fig. 3**). The geographic location of these subjects spanned 5 countries (USA, China, Denmark, Austria, and Luxembourg). *DsrAB* was detected in 30.6% of subjects: 84.6% of SSRM-positive subjects had SSRM detected from the Desulfovibrionaceae family and 21.4% had SSRM from the Firmicutes group. This is contrasted to a previous large-scale study indicating a SSRM presence in only 17.4% of subjects based on *dsrAB* presence in MAGs (Wolf et al., 2022). This difference may be attributed to the different metagenomic datasets and therefore subjects evaluated, or their use of MAGs that may not retain the less-abundant SSRM.

**Figure 3.**
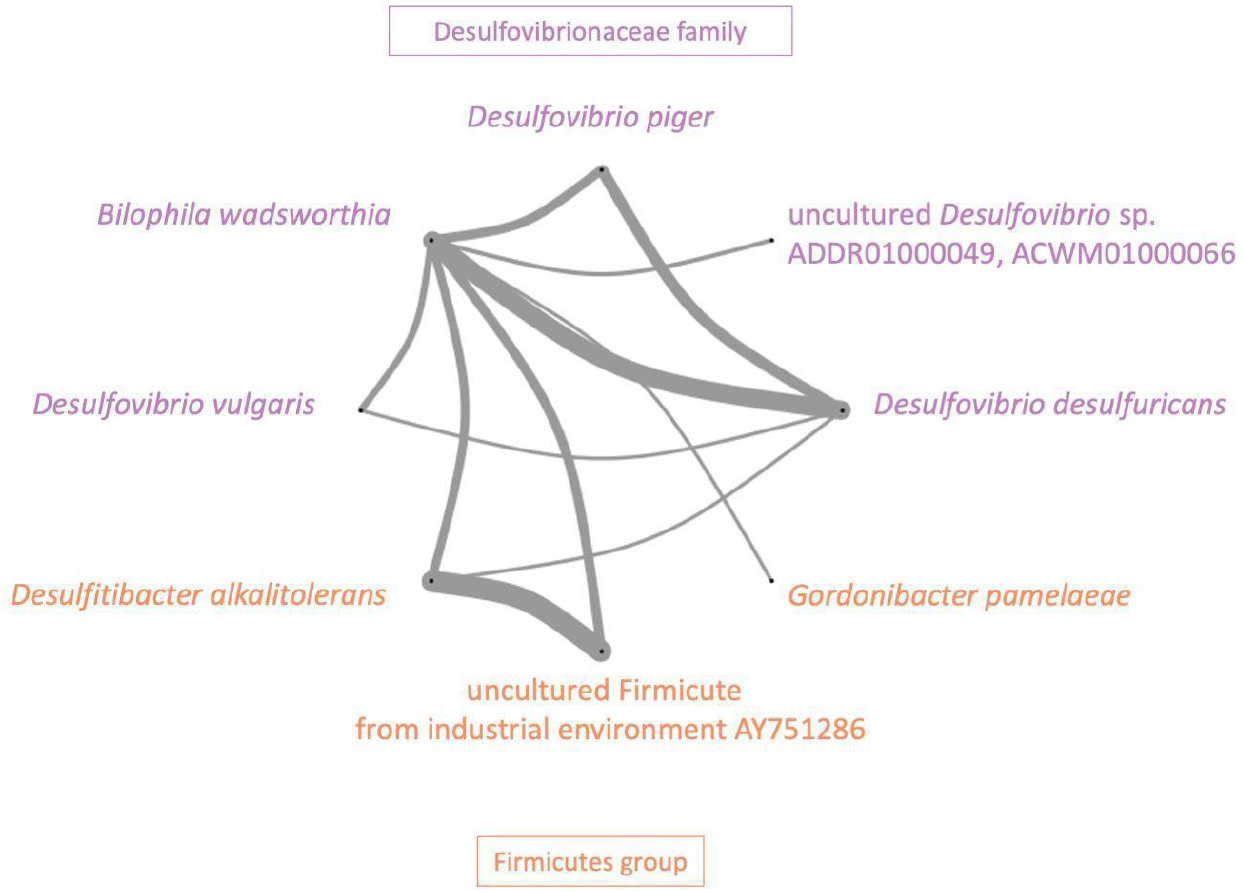
Co-occurrence network of SSRM clusters. SSRM clusters present in at least 20 subjects were included in this analysis. Edge width is proportional to the number of observations (*n* subjects with both clusters present). Representative SSRM are colored by superclusters, according to Figure 1A.

Genera from the Desulfovibrionaceae family, including *Bilophila* and *Desulfovibrio*, represented the most prevalent SSRM (**Fig. 1B**). Of all 4,228 subjects and the 1,294 SSRM-positive subjects, 18.3% and 59.7% contained hit sequences in the *Bilophila wadsworthia* cluster. Although 84.6% of SSRM-positive subjects had members of the Desulfovibrionaceae family, this value drops to 38.9% when excluding the *B. wadsworthia* cluster. Following *Bilophila*, the *Desulfovibrio desulfuricans* and the *Desulfovibrio piger* clusters were of the highest prevalence at 20.8% and 16.9% of SSRM-positive subjects. The remaining Desulfovibrionaceae clusters were individually present in less than 5% of SSRM-positive subjects. SSRM from the Firmicutes group were found in 10.8% of all and 21.4% of SSRM-positive subjects. Notably, the most prevalent reference cluster in this group was represented by an uncultured Firmicute from an industrial environment (AY751286 cluster, **Fig. 1B**) which was found in 12.8% of SSRM-positive subjects. 11.8% of SSRM-positive subjects contained sequences from the *Desulfitibacter alkalitolerans* cluster and 6.49% contained sequences from the *Gordonibacter pamelaeae* cluster.

Three hit sequences were most closely related to *Archaeoglobus sulfaticallidus*, a thermophilic archaea. Further evaluation of these samples could not verify the presence of *A. sulfaticallidus* or a close relative in the respective metagenomes (**Supplementary Information**).

### SSRM prevalence and correlation with host age, sex

Next, we analyzed the association of SSRM prevalence with host characteristics. There was a strong association between prevalence and age (point-biserial correlation 0.32, p-value = 3. 82 * 10^−23^). For subjects with age-specific data, the median age of subjects without SSRM was 23 years (interquartile range 1, 46), which was significantly lower than subjects with SSRM (median 42 years, interquartile range 28, 57; **Fig. 2A**). Given the multimodal distribution of age in subjects without SSRM and that the gut microbiota in young children converges to an adult-like configuration by 3 years of age (Yatsunenko et al., 2012), we further analyzed the difference in SSRM presence between subjects under 3 years old and subjects equal to or above 3 years old. A significantly lower percentage of subjects <3 years old were positive for SSRM compared to the 3 years and older group—4.58% and 38.3% respectively (Fisher’s exact test, *p*-value = 2. 20 · 10^−32^). In contrast, no association was observed between SSRM prevalence and sex (**Fig. 2B**).

### SSRM co-occurrence in the gut

To determine the distribution of different SSRM across subjects we performed a co-occurrence analysis of the 8 SSRM clusters that were each identified in 20 or more subjects. The resulting network consisted of 582 instances of co-occurrence along 11 edges (**Fig. 3**). The *wadsworthia* cluster had the highest cumulative degree of co-occurrence; it was connected to 7 other clusters and was included in 60.3% of all instances of co-occurrence. In contrast, the highest weighted edge was observed between the *D. alkalitolerans* and the uncultured Firmicute AY751286 clusters, which was found in 123 subjects and represented the highest degree of co-occurrence. The second highest degree of co-occurrence was between the *B. wadsworthia* and the *D. desulfuricans* clusters, which were found together in 109 subjects. The network further suggested the existence of two dominant patterns of co-occurrence among groups of three SSRM clusters (visible as prominent triangles in the edges in **Fig. 3**.). Both patterns included the *B. wadsworthia* cluster. We further confirmed that all three clusters were co-occurring within the same subject. The triad of *B. wadsworthia, D. alkalitolerans*, and the uncultured Firmicute AY751286 was present in 11.8% of SSRM-positive subjects while the *B. wadsworthia, D. piger*, and *D. desulfuricans* triad was present in 6.2% of SSRM-positive subjects.

## Discussion

Our work identifies the full scope of SSRM diversity in the human gut by utilizing publicly available, partially assembled metagenomic data, allowing us to validate the prevalence of widely recognized SSRM and detect less prevalent species. We support previous reports of *Bilophila wadsworthia* and other Desulfovibrionaceae clusters as the most prevalent SSRM in the human gut (Fite et al., 2004; Jia et al., 2012; Loubinoux et al., 2002; Rey et al., 2013; Wolf et al., 2022) and suggest a few members of the Firmicutes SSRM group (the uncultured Firmicute AY751286, *Desulfitibacter alkalitolerans*, and *Gordonibacter pamelaeae*) as possible key constituents of the gut-SSRM community. Co-occurrence analysis suggests different gut environments can support different groups of co-occurring SSRM. Co-occurrence within, but not between, the Firmicutes group and Desulfovibrionaceae may be explained by metabolic differences in electron acceptor requirements. In addition, *B. wadsworthia* potentially exploits a unique niche, either metabolically by using a unique substrate (e.g. taurine) or spatially by residing in the small intestine, where it is not in direct competition with other colonic SSRM. Furthermore, in contrast to previous studies (Figliuolo et al., 2018; Fite et al., 2004; Jia et al., 2012; Loubinoux et al., 2002), we present a population-level assessment of the prevalence of SSRM in the human gut as our data are not limited to case-control studies evaluating specific disease states. In the primarily industrialized countries represented in our study, SSRM are present in very few children under 3 years old, but are present in the gut microbiome of nearly 40% of adults.

Our approach using partially assembled metagenomic sequencing data allowed for the evaluation of prevalence of SSRM across subjects, but the datasets did not include raw sequencing data, information on total sequencing depth, and contained sparse metadata. This precluded mapping raw reads to obtain abundance estimates of any of the observed *dsrAB* sequences. We were also unable to draw definitive conclusions about the absence of SSRM from subjects in which *dsrAB* was not detected, as there was not a standardized sequencing depth across studies included. As such, the SSRM prevalence we reported is a conservative estimate and could possibly be an underestimation. The paucity of available metadata limited our analysis on the relationship between SSRM and host characteristics. Thus, we echo the importance of standardized reporting criteria (e.g. minimal information standards published by the Genomic Standards Consortium) and web interfaces such as METAGENOTE (Quiñones et al., 2020; Yilmaz et al., 2011), which attempt to ensure genomic sequencing datasets are annotated with a minimum set of host characteristics to maximize utility of data in public databases.

Despite these limitations, we identified the top three most prevalent SSRM clusters were *B. wadsworthia, D. desulfuricans*, and *D. piger*, consistent with previous studies on highly abundant SSRM in the human gut (Y. Feng et al., 2017; Jia et al., 2012; Wolf et al., 2022; Ye et al., 2018). Our findings also confirm the presence of nine additional SSRM clusters from the *Desulfovibrio* genus (Alipour et al., 2016; Balamurugan et al., 2008; Dostal Webster et al., 2019; Fite et al., 2004; Kleessen et al., 2002; Loubinoux et al., 2002; Murros et al., 2021). Our study is the first to report *D. carbinoliphilus* in the human gut and four additional *Desulfovibrio* clusters that have not been reported in any human-associated microbiome.

Until recently, SSRM in the Firmicutes group have been overlooked as members of the gut microbial community potentially responsible for H_2_S production (Wolf et al., 2022). We identified 13 SSRM clusters within the Firmicutes group. The uncultured Firmicute AY51286 and *Desulfitibacter alkalitolerans* were the most prevalent. Neither have been reported in the human gut, but were present in 70-75% as many subjects as the commonly studied *D. piger*. Interestingly, both were originally detected in industrial heating plant biofilms in Denmark (Kjeldsen et al., 2007; Nielsen et al., 2006), but in the present study were detected in over 150 human subjects from the USA, China, Denmark and three other European countries, indicating they are prevalent members of the gut microbiota from industrialized countries globally. Evaluation of the abundance and *in vivo* sulfite reduction capacity by these commonly overlooked, but prevalent, SSRM clusters may reveal their importance for H_2_S production in the human gut.

We discovered prominent co-occurrence patterns in two different triads of SSRM: *B. wadsworthia*/*D. alkalitolerans*/uncultured Firmicute AY751286 (Firmicute triad) and *B. wadsworthia*/*D. piger*/*D. desulfuricans* (Desulfovibrionaceae triad). *Desulfovibrio* species all utilize free sulfate as the terminal electron acceptor in anaerobic respiration (Steger et al., 2002), so gut environments containing these resources may be advantageous to all SSRM from *Desulfovibrio*. The Desulfovibrionaceae triad contains the three most prevalent SSRM detected in our study and therefore may itself be an artifact of the most prevalent SSRM. However, the lack of co-occurrence between the Desulfovibrionaceae SSRM, with the exception of *B. wadsworthia*, and SSRM from the Firmicutes group is notable and suggests a potentially different gut environment. *D. alkalitolerans* and the uncultured Firmicute AY751286 both use elemental sulfur, sulfite, and thiosulfate—but not sulfate— for metabolism (Kjeldsen et al., 2007; Nielsen et al., 2006), suggesting potential differences in the gut environments (e.g. redox state, metabolites presents, etc.) of individuals who harbor one triad over another. In contrast, *B. wadsworthia* can metabolize sulfite from taurine after cleavage from taurocholic acid and has been detected in the human small intestine, where bile acids are present at 10x higher concentration (Shalon et al., 2023), so it is possible that it uniquely utilizes this substrate and/or inhabits a different spatial area of the gut than other SSRM species.

Our study reports the overall prevalence and diversity of SSRM in the human gut including low abundance species, showing that over a third of subjects from industrialized countries contain genes capable of dissimilatory sulfite reduction in their gut microbiome, supporting and extending previous findings (Wolf et al., 2022). We show that SSRM in the gut are limited to two main clades of the phylogenetic tree composed of all known *dsrAB*-containing organisms from diverse environments. Further, our results show that the understudied Firmicutes group adds important diversity to the gut-SSRM community, and is likely present in individuals with a surplus of sulfite, but not sulfate, in their gut environment. Understanding these nuances will assist researchers in the future who aim to understand the negative health effects of microbially-produced H_2_S in the human gut.

## Materials and Methods

### Construction of pHMMs

HMMER (Eddy, 2011; Finn et al., 2011) determines sequence homology by comparing a target sequence, in this case open reading frames derived from assembled contigs, to a profile Hidden Markov Model (pHMM), a statistical model that evaluates the likelihood of mutation at each position of a multiple sequence alignment of reference sequences, weighting regions conserved across all sequences more heavily. A *dsrAB* reference database was constructed containing 1,292 sequences that cover all publicly available dsrAB sequences from NCBI (status August 2013) (Müller et al., 2015). These 1,292 sequences were used as reference sequences to build two amino acid pHMMs using HMMER3’s hmmbuild tool (Finn et al., 2011). One pHMM was built from bacterial-type *dsrAB* sequences and another from archaeal-type *dsrAB* sequences.

### Determination of score cutoff

34,408 nitrite and sulfite reductase 4Fe-4S domain sequences were downloaded from UniProt on December 8th, 2020 (PF01077). Sequences from this database were identified as *dsrAB* using appropriate keywords (“dsr”, “dissimilatory sulfite reductase”) in the sequence descriptions. A HMMER search was performed with the constructed *dsrAB* pHMMs against this database with default reporting parameters using the hmmsearch tool (Finn et al., 2011). A bit score cutoff of 100 was determined as sufficient in filtering out non-*dsrAB* sequences while retaining *dsrAB* sequences.

### Retrieving human gut metagenomes from NCBI

A list of 72,501 biosample accession numbers from NCBI’s Whole Genome Shotgun database was compiled on February 1st–2nd, 2022, using a combination of search filters specifying human gut metagenomes. For a full list of these search filters, see **Supplementary Table S3**. The associated sequencing data for each biosample was downloaded using the prefetch and vdb-dump commands from the Sequence Read Archive Toolkit (Leinonen et al., 2011) on March 17th–18th, 2022.

### HMMER search for dsrAB

Open reading frames were predicted from assembled contigs obtained from downloaded Biosample datasets and translated to amino acid space using Prodigal (Hyatt et al., 2010). A HMMER search was performed against the translated Biosample datasets with the *dsrAB* pHMMs with default reporting parameters using hmmsearch (Finn et al., 2011). Following the HMMER search, hit sequences were filtered by a bit score of greater than or equal to 100.

### Phylogenetic placement of dsrAB hit sequences

In order to determine the phylogeny of the hit sequences from the HMMER search, phylogenetic placement of the hit sequences onto a reference *dsrAB* tree was performed. The reference *dsrAB* tree was a multifurcating, mixed Maximum Likelihood, Neighbor Joining, and Maximum Parsimony tree constructed from the reference sequence database (Müller et al., 2015). Only unique hit sequences were used for fragment insertion; duplicate hit sequences and shorter hit sequences contained within longer hit sequences were removed. A multiple sequence alignment of the unique hit sequences was made using MAFFT (Katoh & Standley, 2013). Sixteen aligned hit sequences with over 60% gaps were removed from the alignment. The remaining 1,769 unique hit sequences were inserted onto the reference *dsrAB* tree using Random Axelerated Maximum Likelihood (Stamatakis, 2014) with the raxmlHPC command specifying a GAMMA model of rate heterogeneity, a Dayhoff substitution matrix and placement threshold of 0.1, and binary model parameters estimated from the reference *dsrAB* tree.

Following phylogenetic placement, the tree was rooted between the bacterial and archaeal branches. Branch distances between every hit sequence and the closest reference sequence on the tree was calculated using the ETE toolkit (Huerta-Cepas et al., 2016) in order to determine the closest reference relative to each hit sequence. Hit sequences were assigned to the closest reference leaf by branch distance. The 15 hit sequences exceeding a branch distance of 2.5 from the closest reference leaf were manually inspected, identified as oxidative *dsrAB* homologs, and thus removed.

### Clustering

Reference leaves in the *dsrAB* tree were clustered via agglomerative clustering using the tip_glom function from the R package “phyloseq” (McMurdie & Holmes, 2013). A branch height of 0.1 corresponded to >90% nucleotide similarity within clusters. For a list of reference leaves contained within each cluster, see Supplementary Table 1.

### Construction of co-occurrence network

Co-occurrence network was constructed using the R packages “cooccur” (Griffith et al., 2016) and “igraph” (Csárdi & Nepusz, 2006) at the cluster and subject level, only including clusters present in more than 20 subjects.

### Retrieving Biosample host metadata

Metadata information for each Biosample was manually downloaded from NCBI and parsed with an in house script. Biosample hit sequence data was merged on the subject-level by determining individual subjects with the following metadata variables: “host subject id”, “Test person”, “submitted subject id”, “patient_name”, “Patient_name”, “patient”, “individual name”, “individual”, “subject_id”.

Subject age in years was determined by a keyword search for units in days, months, or years under the “host age” variable, and subjects with age information lacking a unit were manually checked for age on NCBI. Data on sample geographic origin was confounded by host age (e.g. some locations only included infant samples), so we did not analyze prevalence by geographic region.

### Statistics

All statistical analyses were performed using R version 4.2.0. Correlation between age and SSRM presence was determined with a point-biserial correlation and one-sided Wilcoxon rank sum test. Correlation between sex and SSRM presence was determined with Fisher’s exact test. The standard p-value cutoff of 0.05 was used to assign statistical significance.

## Data availability

All code (python and R scripts) and datasets are available on GitHub (https://github.com/RebbyAnne/SSRM_pipeline). A list of accession numbers for all Biosamples downloaded from NCBI sequencing read archive and used in this study are included in **Supplementary Table S3** and as a text file on the GitHub site.

## Acknowledgements

We would like to thank Les Dethlefsen, Eitan Yaffe, Dylan Dahan, and Brian Merrill for advice on our methodological approach. We acknowledge the Stanford Research Computing Center for providing computational resources (Sherlock cluster). JAG was supported by a Stanford University School of Medicine Dean’s Fellowship. RAC was supported by the Bio-X Stanford Undergraduate Research Program and a Stanford Vice Provost for Undergraduate Education Major Grant. Funding for the project was also supported by a Stanford Microbiome Seed Grant to JAG and AMS (2015-2016). JC acknowledges support by a Stanford Bio-X Seeding Grant (grant number 10-32), a Center for Pediatric IBD and Celiac Disease Seed Grant (309906), and a Terman fellowship. The funders had no role in study design, data collection and interpretation, or the decision to submit the work for publication.

