## Supplementary Information for "Diversity and prevalence of sulfate- and sulfite-reducing microorganisms in the human gut microbiome"

### Supplementary Text

#### Investigation into presence of *Archaeoglobus sulfaticallidus*

The three biosamples containing these hit sequences for *A. sulfaticallidus* were BLASTed against the *A. sulfaticallidus* genome, resulting in relatively few, very short hits. When those hits in turn were BLASTed against the NCBI nucleotide database, a majority of the most closely related sequences belonged to members of the *Lachnospiraceae*, a Firmicutes family that so far is not known to contain *dsrAB*.

### Supplementary Figures

**A.**

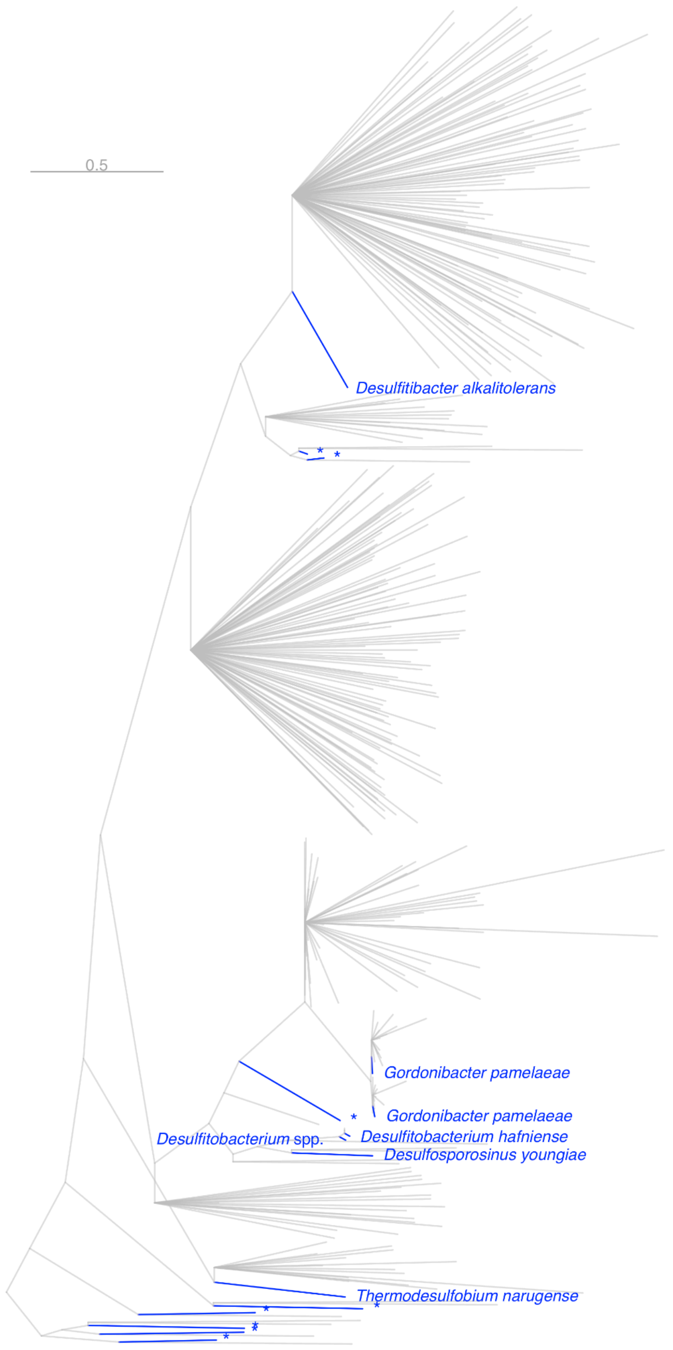

**B.**

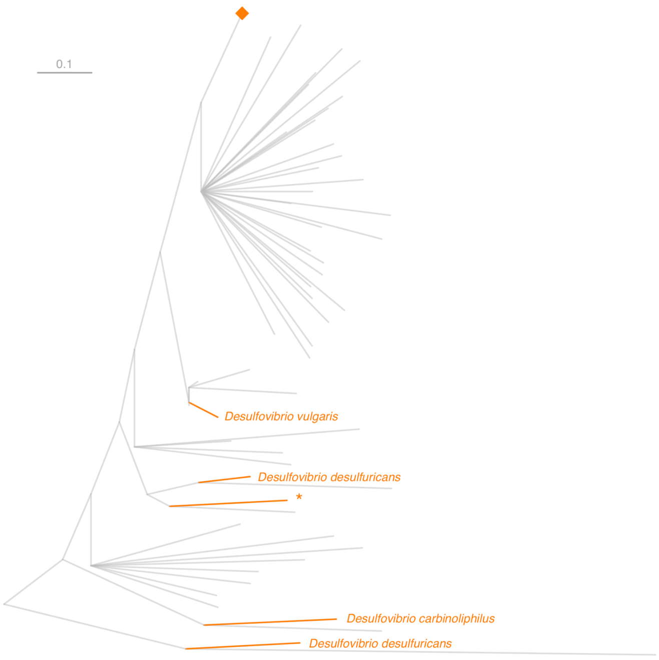
0.1
Desulfovibrio vulgaris
Desulfovibrio desulfuricans
*
Desulfovibrio carbinoliphilus
Desulfovibrio desulfuricans

**C.**

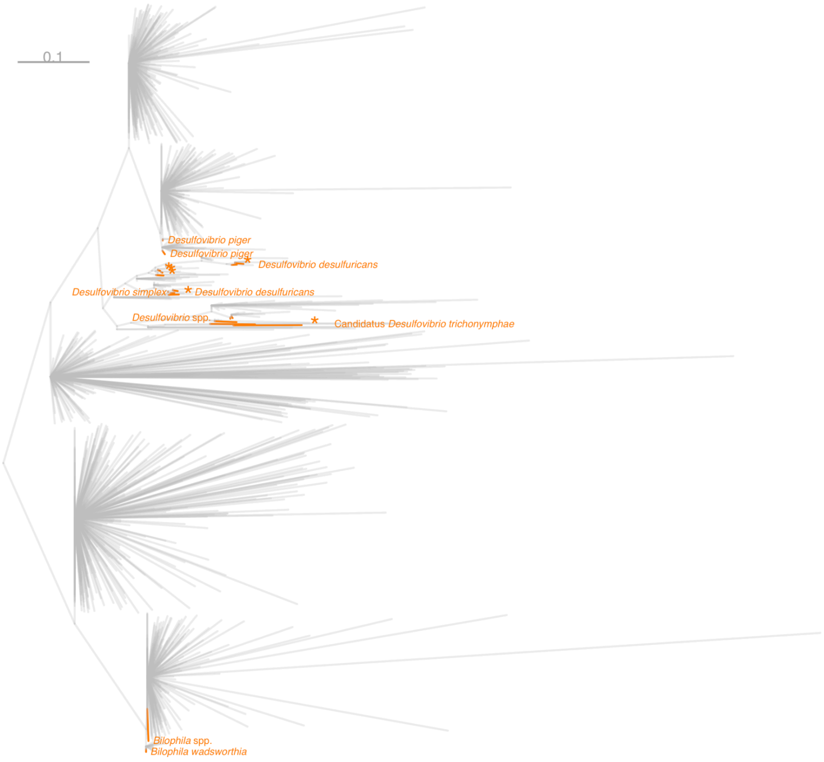
0.1
Desulfovibrio piger
Desulfovibrio piger
Desulfovibrio desulfuricans
Desulfovibrio simplex
Desulfovibrio desulfuricans
Desulfovibrio spp.
Candidatus Desulfovibrio trichonymphae
Bilophila spp.
Bilophila wadsworthia

**Supplementary Figure 1.** Phylogenetic tree of A) Firmicutes group and B,C) Desulfovibrionaceae family. Collapsed clade in B) shown expanded in C). Reference leaves are colored (blue for Firmicutes, orange for Desulfovibrionaceae) and inserted hit sequences are grey. Labels are for closest reference leaves. Unclassified reference leaves marked with asterisk (*).

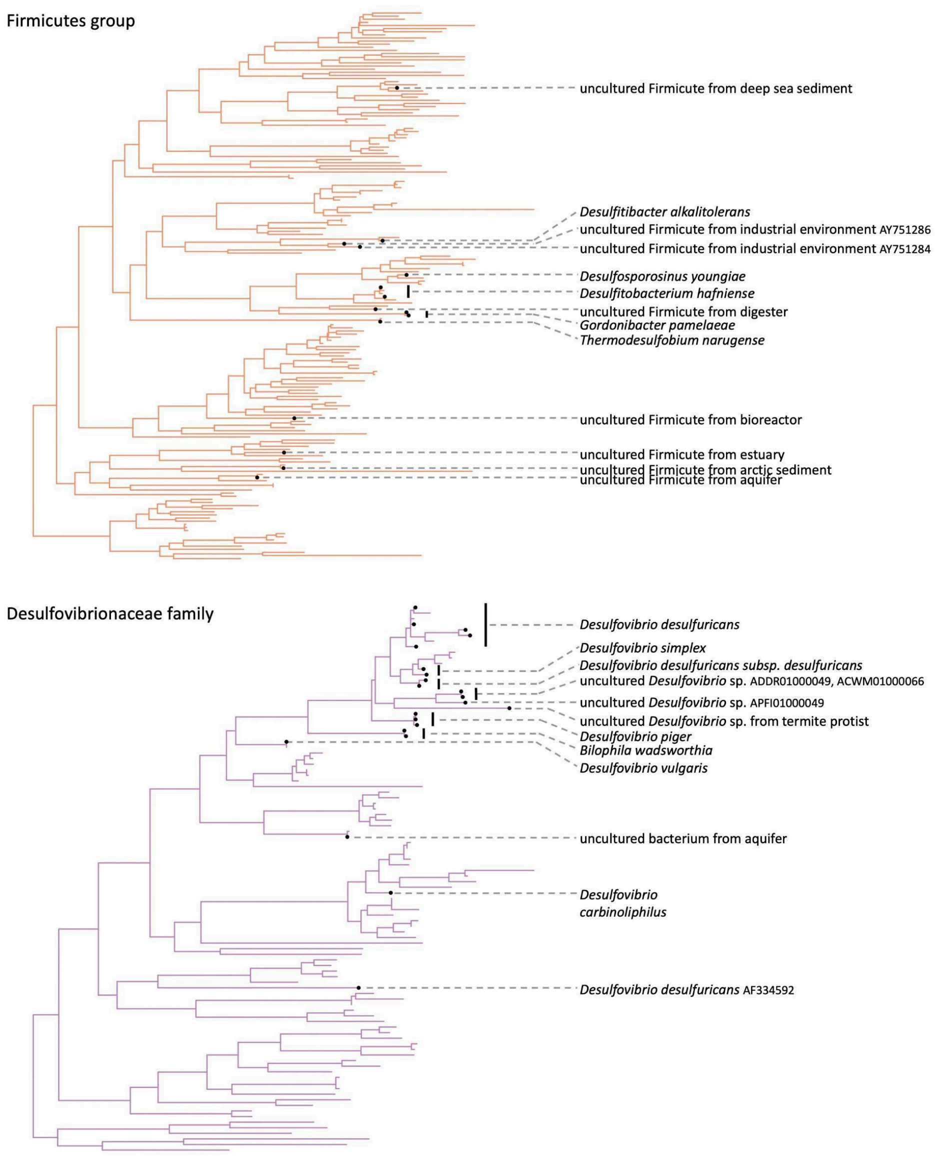


**Supplementary Figure 2.** Reference trees for the Firmicutes group (top, orange) and Desulfovibrionaceae family (bottom, purple), clustered to >90% nucleotide sequence identity. Reference leaves that were closest by branch length to hit sequences are labeled with black points and the cluster representative name.

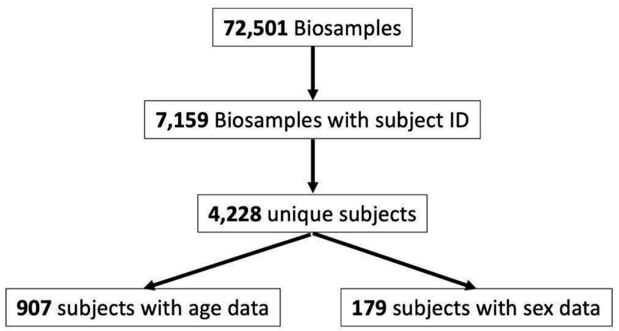


**Supplementary Figure 3.** Flowchart of number of biosamples, subjects, and subject metadata. Some studies were longitudinal, provided individually assembled MAGs as unique biosamples, or had study designs that were not specified and did not include subject identifiers in the uploaded metadata, etc.

**Supplementary Tables**:

**Supplementary Table S1**:
https://github.com/RebbyAnne/SSRM_pipeline/blob/main/config/metadata/NCBI_biosample_project_accessions.csv
List of BioSample IDs and corresponding BioProjects used in this study.

**Supplementary Table S2**:
https://github.com/RebbyAnne/SSRM_pipeline/blob/main/Supplementary_Table_S2_SpeciesLevelClusters.xlsx
Assigned species-level clusters with >90% dsrAB nucleotide sequence identity for

**Supplementary Table S3**:
https://github.com/RebbyAnne/SSRM_pipeline/blob/main/Supplementary_Table_S3_FiltersForWGSDatabase.xlsx
Search filters to specify human gut metagenomes from NCBI's Whole Genome Shotgun database for compiled list of BioSample accession numbers.
